# In vivo Correlation Tensor MRI reveals microscopic kurtosis in the human brain on a clinical 3T scanner

**DOI:** 10.1101/2021.11.02.466950

**Authors:** Lisa Novello, Rafael Neto Henriques, Andrada Ianuş, Thorsten Feiweier, Noam Shemesh, Jorge Jovicich

## Abstract

Diffusion MRI (dMRI) has become one of the most important imaging modalities for noninvasively probing tissue microstructure. Diffusion Kurtosis MRI (DKI) quantifies the degree of non-gaussian diffusion, which in turn has been shown to increase sensitivity towards, *e.g.*, disease and orientation mappings in neural tissue. However, the specificity of DKI is limited as different microstructural sources can contribute to the total diffusional kurtosis, including: variance in diffusion tensor magnitudes (*K*_iso_), variance due to intravoxel diffusion anisotropy (*K*_aniso_), and microscopic kurtosis (*μK*) related to restricted diffusion and/or microstructural disorder. The latter in particular is typically ignored in diffusion MRI signal modeling as it is assumed to be negligible. Recently, Correlation Tensor MRI (CTI) based on Double-Diffusion-Encoding (DDE) was introduced for kurtosis source separation and revealed non negligible *μK* in preclinical imaging. Here, we implemented CTI for the first time on a clinical 3T scanner and investigated the kurtosis sources in healthy subjects. A robust framework for kurtosis source separation in humans is introduced, followed by estimation of the relative importance of *μK* in the healthy brain. Using this clinical CTI approach, we find that *μK* significantly contributes to total diffusional kurtosis both in gray and white matter tissue but, as expected, not in the ventricles. The first *μK* maps of the human brain are presented. We find that the spatial distribution of *μK* provides a unique source of contrast, appearing different from isotropic and anisotropic kurtosis counterparts. We further show that ignoring *μK* - as done by many contemporary methods based on multiple gaussian component approximation for kurtosis source estimation - biases the estimation of other kurtosis sources and, perhaps even worse, compromises their interpretation. Finally, a twofold acceleration of CTI is discussed in the context of potential future clinical applications. We conclude that CTI has much potential for future *in vivo* microstructural characterizations in healthy and pathological tissue.

**Highlights:**

- Correlation Tensor MRI (CTI) was recently proposed to resolve kurtosis sources
- We implemented CTI on a 3T scanner to study kurtosis sources in the human brain
- Isotropic, anisotropic, and microscopic kurtosis sources were successfully resolved
- Microscopic kurtosis (*μK*) significantly contributes to overall kurtosis in human brain
- *μK* provides a novel source of contrast in the human brain in vivo

## Introduction

Mapping microstructural features of the brain, *in vivo* and non-invasively, has been a central endeavor of the MRI community for more than two decades. Diffusion-weighted MRI (dMRI) has played a key role in such microstructural characterizations due to its ability to sensitize the MRI signal towards diffusion-driven displacements, which then “sense” tissue boundaries in the range of around >10 μm (in typical clinical settings). Water molecules are highly abundant in tissues and can traverse several micrometers in a typical MR-relevant diffusion time, making dMRI an excellent indicator of tissue microstructure. At relatively low diffusion weighting, the unidirectional Apparent Diffusion Coefficient (ADC, Stejskal and Tanner, 1965; Le Bihan et al., 1986) and later the rotationally invariant Diffusion Tensor Imaging approaches (DTI, Basser et al., 1994) utilized a Gaussian diffusion framework for quantifying diffusivities, which found numerous applications from stroke detection, to white matter orientation mapping, to characterizing progressive changes in brain tissue due to plasticity (*e.g.* Moseley et al., 1990; Gauvin et al., 2001; Anwander et al., 2007; Roebroeck et al., 2008; Della-Maggiore et al., 2009; Scholz et al., 2009; Blumenfeld-Katzir et al., 2011; McNab et al., 2013a; Benetti et al., 2018; Hasan et al., 2018; Jacobacci et al., 2020, Yon et al., 2020, for a review see *e.g.* Mukherjee, 2005; Assaf and Pasternak, 2008; Johansen-Berg, 2010).

Deviations from Gaussian displacement profiles were described quite early within the framework of q-space MR (Callaghan et al., 1991; Assaf and Cohen, 1998). As high performance magnetic gradients become more readily available in clinical MRI systems, a wider range of diffusion-weighting regimes could be probed, providing evidence for non-gaussian profiles also in the human brain (Mulkern et al., 1999). Since q-space gradients required very high diffusion weighting and suffer from low signal to noise, a general methodology for characterizing diffusion weighted signals at an intermediate diffusion weighting regime was elegantly introduced by Jensen et al., 2005 and Jensen & Helpern, 2010. The ensuing Diffusion Kurtosis Imaging (DKI) methodology is a “signal representation” approach (Novikov et al., 2018; Novikov, 2021, as opposed, *e.g.*, to microstructural models, for instance: Jespersen et al., 2007; Fieremans et al., 2011; Zhang et al., 2012; Kaden et al., 2016; Jespersen, 2018) based on the cumulant expansion of the dMRI signal up to the second order in *b*-value (with *b* = *γ*^2^*δ*^2^*G*^2^(*Δ*-*δ*/3) for ideal rectangular diffusion-sensitizing gradients, where *γ* is the gyromagnetic ratio, *δ* is the gradient pulse duration, *Δ* is separation between the two leading edges of the gradient pulses, and *G* is the gradient pulse’s magnitude). By quantifying the kurtosis in water displacement probability, a new source of contrast for tissue microstructure was introduced (Jensen et al., 2005; Jensen & Helpern, 2010; Wu & Cheung, 2010). DKI has been successfully adopted to study both the healthy and the diseased human brain, providing insights into individual anatomy at high resolution (Mohammadi et al., 2015), development (Huber et al., 2019), aging (Falangola et al., 2008; Henriques, 2018), attention deficit hyperactivity disorder (ADHD, Helpern et al., 2010), neurodegenerative disorders (Arab et al., 2018) such as Alzheimer’s (Gong et al., 2013, Struyfs et al., 2015) and Parkinson’s disease (Wang et al., 2011; Kamagata et al., 2013; Kamagata et al., 2014; Surova et al, 2018), for characterizing brain tumors (Raab et al., 2010; Raja et al., 2016; Falk Delgado et al., 2017; Hempel et al., 2017; Hempel et al., 2018; Lin et al., 2018), and is nowadays implemented in different platforms (*e.g.* Leemans et al., 2009; Tabesh et al., 2011; Tournier et al., 2019; Henriques et al., 2021a).

Since DKI can be thought of as more sensitive to tissue heterogeneity, it was recognized relatively early on that it can provide higher sensitivity than its DTI counterpart (*e.g.* Falangola et al., 2008; Wang et al., 2011; Hui et al., 2012; Zhuo et al., 2012; Fieremans et al., 2013; Steven et al., 2014; Lin et al., 2018). However, at the same time, DKI suffers from a lack of specificity, since many factors can contribute to heterogeneity. At the voxel level, diffusion kurtosis may arise in the presence of mesoscopic effects such as the orientation dispersion of fibers and their specific configuration (Lu et al., 2006; Henriques et al., 2015). At the microscopic level (*i.e.* (sub)cellular level, or more generally at the level of pores or microdomains, where a microdomain can be defined as a uniform sub-voxel segment, Szczepankiewicz et al., 2015), diffusion kurtosis may arise from different sources, including the following ones (Henriques et al., 2020):

1. variance of the eigenvalues of individual diffusion tensors representing tissue microdomains, thus in the presence of *shape* variance (*i.e.* deviations of the shape of pores or microdomains from a sphere), which is referred to as anisotropic kurtosis (*K*_aniso_);
2. variance across the ensemble of all microdomains’ tensor magnitudes, thus in the presence of *size* variance, which is referred to as isotropic kurtosis (*K*_iso_; the subscript “iso” here refers to the isotropic part of the tensor (its magnitude), and does not require the tensor itself to be isotropic);
3. non-Gaussian diffusion within restricting “pores” (restricted diffusion) and/or a combination of restriction and exchange in case components are exchanging, which is referred to as microscopic kurtosis (*μK,* also previously called intra-compartmental kurtosis, *K*_intra_, in Henriques et al., 2020).

Understanding how each of these different sources of tissue heterogeneity affect the kurtosis signal is challenging. DKI, and more generally techniques based on Single Diffusion Encoding sequences (SDE, Shemesh et al., 2016) with moderate *b*-values do not allow to determine the relative contribution of each source without making strong assumptions about tissue properties (Fieremans et al., 2011; Ianuş et al., 2016). Such assumptions, however, may affect the specificity of the derived metrics (Jelescu et al., 2015; Lampinen et al, 2017; Henriques et al., 2019), leading to potential serious errors in the “microstructural” interpretation of the metrics (Jelescu et al., 2016; Novikov et al., 2018).

Nevertheless, the ability to characterize each diffusion kurtosis source has the potential of unraveling important microstructural information that can provide new insights into tissue properties which, in turn, may have clinical impact (Szczepankiewicz et al., 2015; Szczepankiewicz et al., 2016; Nilsson et al., 2020; Alves et al., 2021). For this reason, Multidimensional Diffusion Encoding (MDE, Eriksson et al., 2013; Lasic et al., 2014, Westin et al., 2016 Shemesh et al., 2016; Topgaard, 2017), realized by combining or concatenating multiple diffusion encodings, has been developed to produce richer diffusion weighting paradigms allowing to explore correlations between different spatial dimensions, and, ultimately, to yield more specific tissue contrasts. Among MDE preparations, Double Diffusion Encoding (DDE, Mitra, 1995; Shemesh et al., 2010a; Shemesh et al., 2016, Henriques et al., 2021b) in the long mixing time regime has been used in combination with either trapezoidal or oscillating gradients (Double Oscillating Diffusion Encoding, DODE, Ianuş et al., 2017; Shemesh, 2018). Such applications stemmed from the initial theoretical ground (Mitra, 1995; Cheng & Cory, 1999; Özarslan and Basser, 2008; Özarslan, 2009; Lawrenz et al., 2010; Jespersen & Buhl, 2011; Jespersen, 2012) and the following experimental observations confirming that DDE provides signal differences for collinear and orthogonal diffusion gradients in elongated samples (Cheng & Cory, 1999; Callaghan & Komlosh, 2002) and angular modulations of the signal in the presence pores characterized by eccentricity over the plane sampled by the diffusion preparation (Shemesh et al., 2010a; Shemesh et al., 2010b; Shemesh et al., 2011; Shemesh et al., 2012a). Since then, DDE in the long mixing time regime has been extensively adopted to provide estimations of the microscopic diffusion anisotropy (*μA*), a measure related to *K*_aniso_ by the relationship *K*_aniso_ = 2(*μA*^2^/*D*^2^), with *D* being the mean diffusivity, *μA^2^* ≡ 3/5*var*(*λ*_i_), and *λ*_i_ being the eigenvalues of individual diffusion tensors representing tissue microdomains (Jespersen et al., 2013; Lasic et al., 2014; Shemesh et al., 2016; Ianuş et al., 2018; Henriques et al., 2021b). The *μA* measure, and its normalized version *μFA*, thus report on the anisotropy of the tissue at the pore or microdomain length scale (*i.e.* on its eccentricity), without the confounding effect of orientation dispersion at the macroscopic (*i.e.* at the voxel) level. DDE-derived *μA* has been investigated and successfully observed in preclinical systems in both white and grey matter: in fixed grey matter (Komlosh et al., 2007), pig spinal cord (Komlosh et al., 2008), pig optic nerve and cortical grey matter, with phantoms mimicking their respective microstructures (Shemesh et al., 2010b; Shemesh et al., 2011; Shemesh and Cohen, 2011; Shemesh et al., 2012a), in both *in vivo* and *ex vivo* rat brain (Shemesh et al., 2012b, Ianuş et al., 2018; Kerkelä et al., 2020), in *ex vivo* monkey brain (Jespersen et al., 2013), in rat spinal cord injury (Budde et al., 2017), and in rat spinal cord *ex vivo* (Shemesh, 2018). In clinical systems DDE-derived *μA* has been reported in pig spinal cord (Lawrenz and Finsterbusch, 2011), and successfully mapped in the *in vivo* healthy human brain white matter (Lawrenz and Finsterbusch, 2013; Lawrenz and Finsterbusch, 2015), cortical grey matter (Lawrenz and Finsterbusch, 2019), in the study of aging (Lawrenz et al., 2016), and in clinical applications, where it showed increased specificity compared to DTI-derived FA in multiple sclerosis lesions (Yang et al., 2018) and has been investigated in Parkinson’s disease (Kamiya et al., 2020).

In addition to DDE preparations, alternative MDE pulse sequences have been adopted in parallel with the aim of resolving the kurtosis sources: previous studies showed that, under the strict Multiple Gaussian Component assumption (MGC, Henriques et al., 2020; Henriques et al., 2021c), tensor-valued information of MDE sequences can decompose *K*_t_ as the sum of *K*_aniso_ and *K*_iso_ (Szczepankiewicz et al., 2015, Szczepankiewicz et al., 2019a, Topgaard, 2019). While successful applications of such techniques demonstrated great promise for clinical usefulness (Szczepankiewicz et al., 2015; Szczepankiewicz et al., 2016; Andersen et al., 2020) and implementation (Szczepankiewicz et al., 2019a; Nilsson et al., 2020), a drawback of these so-called “tensor-valued” approaches is that they assume that all underlying diffusion propagators within the voxel can be approximated as multiple Gaussian components (Henriques et al., 2020; Henriques et al., 2021c). However, the validity of these assumptions has recently been called into question (Jespersen et al., 2019; Henriques et al., 2020; Henriques et al., 2021c), mainly due to the following two reasons:

1. observations of time dependent diffusion and kurtosis, a hallmark of diffusion non-Gaussianity in at least one tissue compartment (Novikov et al., 2014; Fieremans et al., 2016; Novikov et al., 2019; Lee et al., 2020a), both in preclinical systems (*in vivo*: Does et al., 2003; Kunz et al., 2013; *ex vivo*: Assaf and Cohen, 1998; Aggarwal et al., 2012; Portnoy et al., 2013; Ianuş et al., 2018; Jespersen et al., 2018) and in clinical settings *in vivo* (Horsfield et al., 1994; Baron and Beaulieu, 2014; Van et al., 2014; Fieremans et al., 2016; Lee et al., 2018; Grussu et al., 2019; Arbabi et al., 2020; Lee et al., 2020a; Lee et al., 2020b), with anisotropic tissues generally presenting residual orientational time-dependency in isotropic encodings (Jespersen et al., 2019);
2. the mapping of non-vanishing positive *μK* values in phantoms and both *in vivo* and *ex vivo* mice at different field strengths (Paulsen et al., 2015; Henriques et al., 2020; Henriques et al., 2021c; Alves et al., 2021).

To address these concerns and resolve kurtosis sources without explicitly or implicitly relying on the MGC assumption, a new DDE strategy has been recently proposed and evaluated in preclinical systems, namely the Correlation Tensor Imaging, or CTI (Henriques et al., 2020, Henriques et al., 2021c). In particular, CTI enables the simultaneous estimation of *K*_iso_, *K*_aniso_, and *μK*, by relying on the acquisition of four different DDE sets (*i.e.* combinations of gradient amplitude and direction) and on the cumulant expansion of DDE signals. So far, the CTI methodology has been tested in both *in vivo* and *ex vivo* rodent brains (Henriques et al., 2020, Henriques et al., 2021c; Alves et al., 2021). These studies revealed the expected contrasts for *K*_aniso_, the dominant kurtosis source in white matter, and for *K*_iso_, which highlights areas with large dispersion of microdomains’ mean diffusivities such as regions with partial volume effects. Crucially, the *in vivo* CTI data demonstrated for the first time evidence of a positive non-vanishing *μK* both in white and grey matter, with *μK* being the dominant kurtosis source in grey matter (Henriques et al., 2021c). However, the CTI framework has not yet been investigated in the human brain. Together with the observation for biases in *K*_aniso_ and *K*_iso_ when not accounting for *μK* in mice (Henriques et al., 2021c), this body of evidence prompts the translation of the CTI methodology in the clinical setting, building also on previous successful applications of DDE sequences in the study of the living human brain.

In this work, we set out to develop CTI for humans using a clinical 3T system and investigate the kurtosis sources emerging from a cohort of healthy subjects. In particular, the goals of this study are i) to evaluate the feasibility of acquiring DDE data adhering to the CTI methodology proposed in Henriques et al., 2021c, in a clinical 3T system; ii) to estimate the kurtosis sources in brain tissue derived by the CTI framework from healthy adult volunteers, particularly with respect to evaluating evidence for the *μK* component in humans; and iii) to investigate the possible implications of ignoring the *μK* component, as currently established in alternative approaches aiming to resolve kurtosis sources.

## Materials and Methods

### Correlation Tensor Imaging Theory

In the long mixing time regime, the powder average of DDE signals is equivalent to (Henriques et al., 2021c):

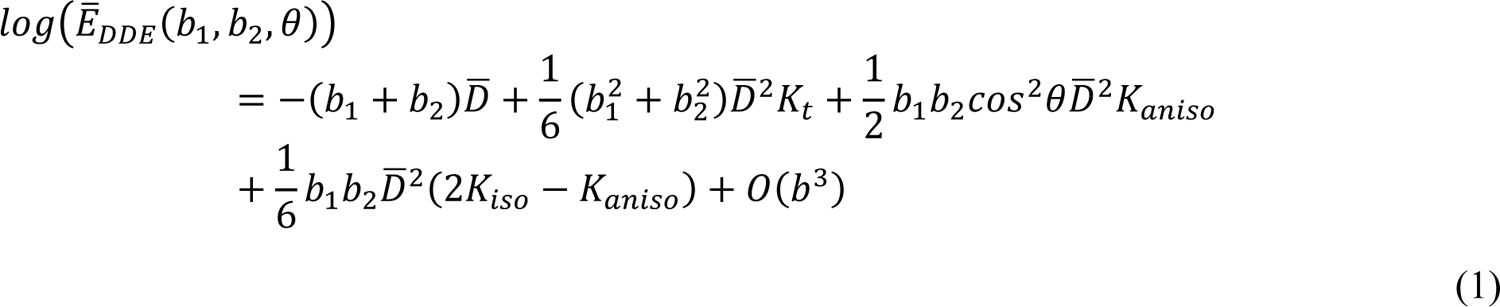

with *b_1_* and *b_2_* being the *b*-values of the first and of the second pair of diffusion gradients, respectively, *θ* being the angle between the directions of the first and the second pair of diffusion gradients, ^θ^ being the mean diffusivity, and *K*_t_ being the total kurtosis of the powder averaged signal. By estimating *K*_t_, *K*_aniso_, and *K*_iso_, then *μK* can be extracted by subtraction as follows (Henriques et al., 2021c):

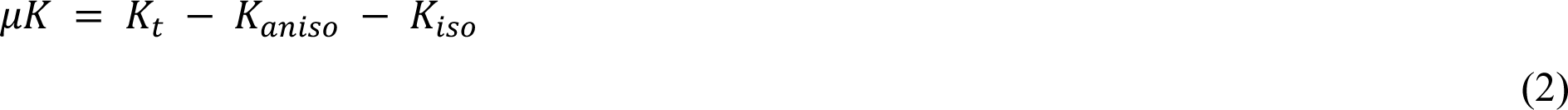

Thus, while *K*_t_ can be accessed by conventional SDE experiments with at least two non-zero *b*-values, its decomposition in its sources requires the use of DDE preparations. Importantly, as described in Henriques et al., 2021c, the difference between the logarithm of powder averaged signals from a SDE-like set with *b*-value *b*_a_ and a parallel DDE set with the

*b-* value of each pair of gradients being *b*_a_/2 can be used to directly access *μK* as follows:

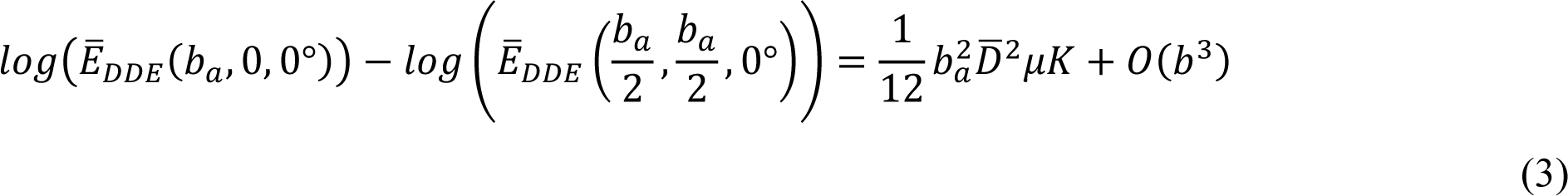

and, in addition, *K*_aniso_ can be directly estimated from the difference between the logarithm of parallel and perpendicular sets (Jespersen et al., 2013, Ianuş et al., 2018):

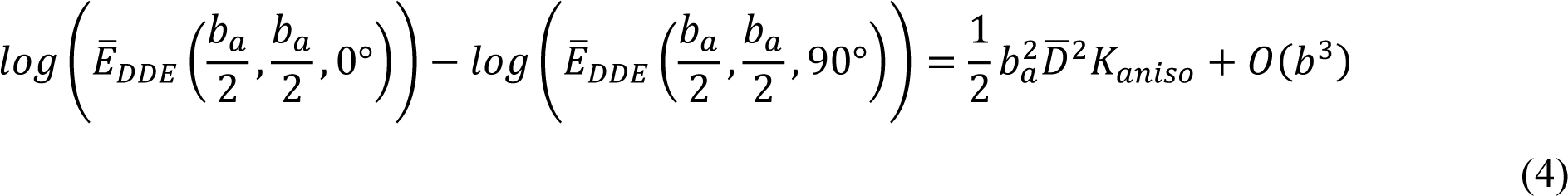

with the higher order terms *O(b^3^)* ignored in the current analysis.

### Participants

Ten participants (mean age ± one standard deviation (SD): 28.9 ± 6.0 years, six males) gave informed consent to participate in this study approved by the Research Ethics Committee of the University of Trento. At the time of MRI scans, all participants were healthy and had no medical history of neurological or psychiatric disorders.

### MRI acquisition

Data were acquired at the Center for Mind/Brain Sciences of the University of Trento, Italy, on a 3T clinical MR scanner (MAGNETOM Prisma, Siemens Healthcare, Erlangen, Germany), with a 64-channel head-neck RF receive coil. Head motion during the acquisition was limited by the use of foam paddings optimized for head coils inside the coil. Anatomical T1-weighted (T1w) Multi-Echo MPRAGE (ME-MPRAGE, van der Kouwe et al., 2008) images were acquired with the following parameters: TE_1_/TE_2_/TE_3_/TE_4_ = 1.69/3.55/5.41/7.27 ms, TR = 2530 ms, TI = 1100 ms, flip angle: 7°, 1 mm-isotropic resolution, matrix size: 256×256.

For the diffusion-weighted data acquisition, a prototype double-spin-echo DDE sequence was used (Fig. 1A). A double-spin-echo sequence was chosen to mitigate concomitant gradient effects (Callaghan and Komlosh, 2002; Szczepankiewicz et al., 2019b). The DDE sequence imaging parameters were: TR = 5600 ms, TE = 127 ms, matrix size: 84×84, slice thickness: 2.5 mm, no slice gap, spatial resolution: 2.5 mm-isotropic, 60 axial slices allowing full brain coverage, partial Fourier factor: 6/8, GRAPPA/SMS factors: 2/4. Four sets of DDE images were acquired according to the optimization described in Henriques et al., 2021c (Fig. 1B):

- DDE set #1: *b*_1_ = 1000, *b*_2_ = 0, with *b*_t_ = 1000 s/mm^2^;
- DDE set #2: *b*_1_ = 2000, *b*_2_ = 0, with *b*_t_ = 2000 s/mm^2^;
- DDE set #3: *b*_1_ = 1000, *b*_2_ = 1000, with *b*_t_ = 2000 s/mm^2^ and parallel directions for the first and the second pairs of diffusion gradients (*θ* = 0°);
- DDE set #4: *b*_1_ = 1000, *b*_2_ = 1000, with *b*_t_ = 2000 s/mm^2^ and perpendicular directions for the first and the second pairs of diffusion gradients (*θ* = 90°);

**Fig. 1.**
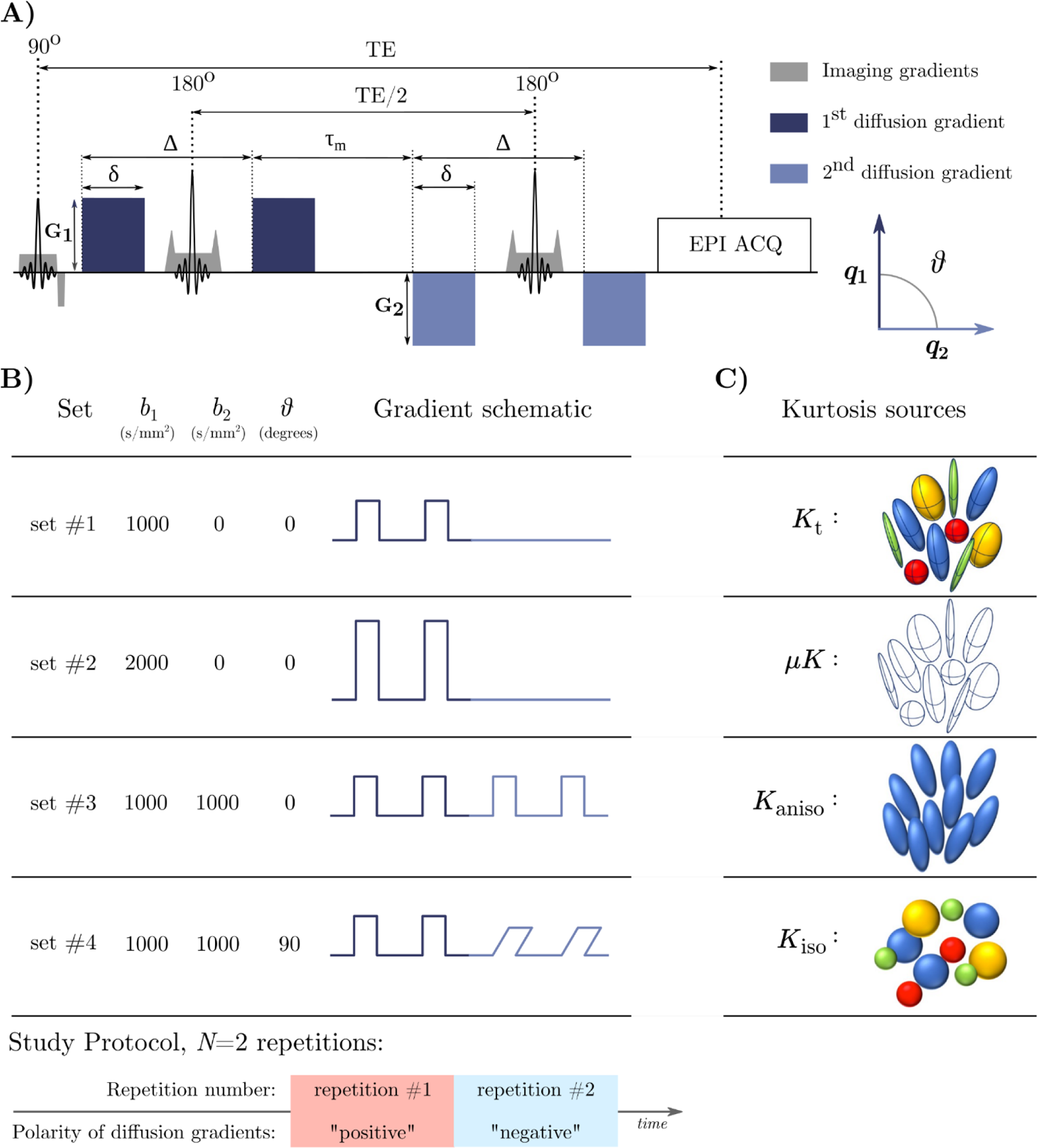
CTI methodology. **A)** DDE sequence adopted in this study. Abbreviations: TE: Echo Time; EPI ACQ: Echo Planar Imaging acquisition module. **B)** CTI acquisition scheme. DDE set #1 and DDE set #2 mimic conventional SDE sequences. The additional DDE set #3 and DDE set #4 are respectively parallel and perpendicular DDE preparations, and they allow to disentangle the total Kurtosis (*K*t) in its Kurtosis sources. **C)** Kurtosis sources. *K*t can be decomposed in: i) the microscopic kurtosis (*μK*), arising from non-Gaussianity induced by restricted time-dependent diffusion, structural disorder such as variations in compartmental cross-sectional area, or exchange effects, which can also be directly accessed through the combination of DDE set #2 and DDE set #3 (see eq. 3); ii) the anisotropic Kurtosis (*K*aniso), arising from the variance in the eigenvalues describing the microdomain diffusion tensors, *i.e.* the shape variance, which can also be directly estimated through the combination of DDE set #3 and DDE set #4 (see eq. 4); and iii) the isotropic Kurtosis (*K*iso), arising from the variance of the microdomain diffusion tensors’ mean diffusivity; factoring out exchange, *K*t = *K*aniso + *K*iso + *μK* (Henriques et al., 2020; Henriques et al., 2021c). The acquisition was repeated twice in the same session without repositioning, with opposite polarities of all the diffusion gradients, allowing to correct for any effects of cross-terms with imaging gradients (Neeman et al., 1991; Lawrenz and Finsterbusch, 2011; Lawrenz and Finsterbusch, 2013; Ianuş et al., 2018).

With *b*_t_ being the total *b*-value, *i.e.* including both the first and the second pairs of diffusion gradients. For all DDE sets, δ/Δ/τ_M_ = 15.8/31.8/32.3 ms. Sixty diffusion-weighted volumes were acquired per each DDE set: for sets #1, #2, and #3 directions from a 3-dimensional 60-point spherical 10-design were used (Hardin & Sloane, 1996), while for set #4, perpendicular directions from the 5-design in Jespersen et al., 2013 were used. For the phantom and all participants, one volume without diffusion gradients (hereafter referred to as “*b*=0” volume) was added every 12 diffusion-weighted volumes, and two additional *b*=0 volumes were acquired at the beginning of set #2 and #4 (except for sub-01 and sub-02, where two b=0 volumes were included per each DDE set and TR = 5500 ms was used). Finally, two *b*=0 volumes with reversed phase encoding were acquired to correct the susceptibility-induced geometric distortions (see *Image processing* section). DDE sets were acquired in the following temporal order to guarantee the possibility of directly estimating *μK* and *K*_aniso_ in case of uncooperative participants leading to the data acquisition interruptions (see eq. 3 and 4): DDE set #2, DDE set #3, DDE set #4, DDE set #1. Two repetitions of the above listed sets were acquired, each with opposite polarity of all diffusion gradients (hereafter referred to as “positive” and “negative”, Fig. 1B, bottom) to reduce potential cross-term effects with imaging gradients (Neeman et al., 1991; Lawrenz and Finsterbusch, 2011; Ianuş et al., 2018). The total acquisition time for the two repetitions was around 52 minutes.

To check for the fulfillment of the long mixing time regime assumption (Shemesh et al., 2012b; Henriques et al., 2020), twelve parallel directions were selected from the gradient scheme for DDE set #3, and the polarity of the direction of the second pair of diffusion gradients was reversed to yield antiparallel gradients (Shemesh et al., 2012b; Henriques et al., 2020). Images corresponding to this subset of parallel and antiparallel gradient directions were acquired twice, each with opposite polarity of the diffusion gradients, followed by ten additional *b*=0 volumes acquired for assessing the temporal Standard Deviation (tSD) of this series and relate it to the signal difference between parallel and antiparallel images for all participants (except for sub-01 and sub-02).

To verify the absence of systematic imaging artifacts, the above described protocol was additionally acquired on a brain-sized spherical isotropic phantom at the room temperature of 22°C, filled with water doped with Nickel Sulphate (NiSO_4_(H_2_O)_6_).

### Image processing

*Diffusion-weighted images*: dMRI data were denoised with the Marchenko-Pastur PCA denoising in MRtrix v. 3.0.2 (Veraart et al., 2016), and derived noise maps were visually inspected along with denoising residuals. Data were then corrected for Gibbs ringing (Kellner et al., 2016), and the susceptibility-induced field was estimated using *b*=0 volumes with opposite phase encoding in FSL’s *topup* (Andersson et al., 2003). Data corresponding to each set was then corrected for eddy currents and head motion in FSL’s *eddy* (Andersson & Sotiropoulos, 2016), by using the first acquired *b*=0 volume as reference volume for all sets, in order to get all the images aligned to the same reference volume. For DDE sets where the amplitude of the second pair of diffusion gradients was not zero (DDE set #3, DDE set #4), the direction of the second pair of diffusion gradients was used in *eddy*, motivated by the observation for a much smaller impact of the first pair of diffusion gradients on eddy currents artifacts (Mueller et al., 2017), and as reported in Yang et al., 2018 and in Fan et al., 2020. Summary metrics of the quality of each participant’s images were then computed with the *eddy qc* framework (Bastiani et al. 2019). Diffusion-weighted volumes were then concatenated according to their temporal order and corrected for signal drift (Vos et al., 2017) and bias field (Tustison et al., 2010). Phantom images underwent the same pre-processing pipeline adopted for human data. Before computing each participant’s (*i.e.* individual) CTI maps, data were smoothed with a 3D Gaussian filter (using a Gaussian kernel with FWHM = 1.25, scipy v. 1.6.1), and pairs of images corresponding to opposite polarities of the diffusion gradients were geometrically averaged to compensate for effects associated with potential imaging cross-terms (Neeman et al., 1991; Lawrenz and Finsterbusch, 2011; Lawrenz and Finsterbusch, 2013; Ianuş et al., 2018). Images were then directionally averaged in order to get four powder average (p.a.) images, one per each set, which were finally normalized (hereafter referred to as p.a._norm_) by the mean of the *b*=0 volumes included across all the acquired DDE sets (Fig. 2A). The tSD across these *b*=0 volumes was computed and used to estimate a SNR map as the ratio of the mean(b=0) and the tSD(b=0) (Fig. 2B).

**Fig. 2.**
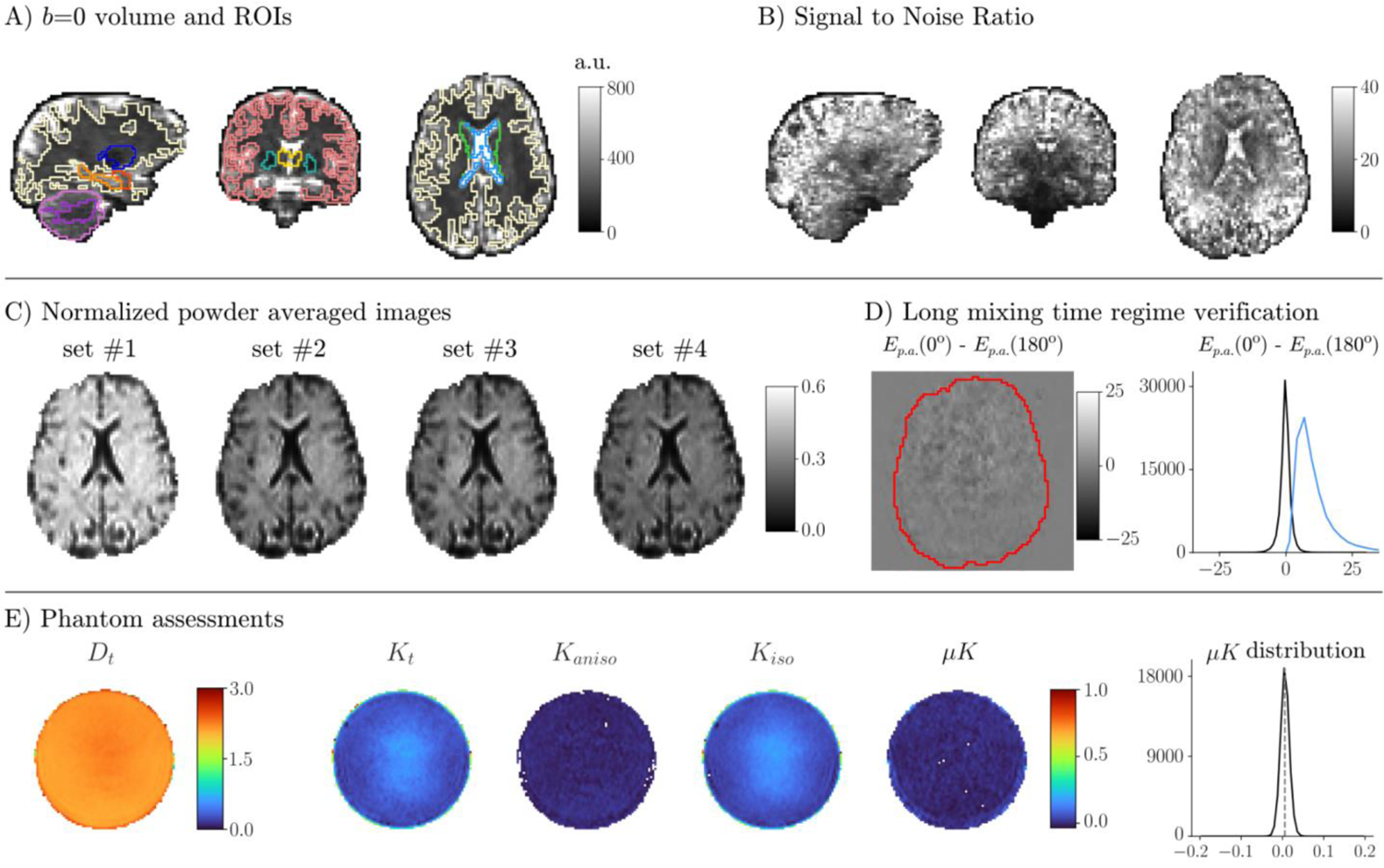
Data quality. All brain images come from the same slices of a representative subject. **A)** *b*=0 volume with overlaid borders of Regions of Interest (ROIs) used in this study. Lateral view: yellow: white matter; blue: putamen; orange: hippocampus; red: amygdala; light violet: cerebellar cortex; violet: cerebellar white matter. Coronal view: pink: cerebral cortex; yellow: thalamus; green: pallidum. Axial view: yellow: white matter; green: caudate; light blue: lateral ventricles. **B)** Lateral, coronal, and axial view of the Signal to Noise Ratio (SNR) map. **C)** Powder (*i.e.* directionally) averaged images per each set normalized by the mean *b*=0 volume (p.a._norm_). **D)** Left: map of the difference between parallel and antiparallel p.a. images at *b*t=2000 s/mm^2^ with the brain border marked in red. Right: distribution of signal differences between the parallel and antiparallel p.a. images (black), with the distribution of the temporal standard deviation values across *b*=0 shown in light blue. **E)** CTI maps derived from the phantom experiment, and distribution of *μK* values within the phantom. Human data underwent the preprocessing pipeline described in the Materials and Methods section, and are shown before the application of the Gaussian smoothing.

*Anatomical images*: T1w images resulting from the root mean squared combination of the four ME-MPRAGE echoes were parcellated with FreeSurfer’s v. 7.1 *recon-all* pipeline. The following Regions Of Interest (ROIs) were considered among the areas included in the FreeSurfer’s parcellation for the quantitative analysis of each kurtosis source: Cerebro-Spinal Fluid (CSF) in lateral ventricles, cerebral White Matter (WM), cerebellar WM (WM_CBM_), Grey Matter (GM), cerebellar Grey Matter (GM_CBM_), Amygdala (AMG), Caudate (Cd), Hippocampus (HPC), Globus Pallidus (GP), Putamen (PU), and Thalamus (TH). Linear registrations between individual skull-stripped T1w images and the first pre-processed non-smoothed *b*=0 volume were then computed in ANTs (Avants et al., 2011) and registration matrices were applied to the above listed ROIs in order to get them aligned with the CTI maps.

*Registration to MNI space and MNI-space CTI analysis*: In order to obtain the first, high SNR group average templates of *K*_aniso_, *K*_iso_, and *μK* in the healthy human brain, CTI was fitted on across-subject averaged p.a._norm_ signals warped to the MNI space. This approach was preferred rather than computing the average of individual CTI maps to avoid the propagation of noise in the individual maps to the final template.

To normalize each participant’s p.a._norm_ images to the MNI space, a non-linear registration between each participant’s skull-stripped T1w image and the MNI 2 mm-isotropic T1w brain atlas was computed. Finally, the resulting warp fields were applied to each p.a._norm_ image in combination with the linear registration matrices previously computed to align the diffusion with the anatomical data (see *Anatomical images* Section), to get all the p.a._norm_ images warped to the MNI space. All registration processes were carried out in ANTs and were visually inspected for their accuracy. No Gaussian smoothing was applied on data that were normalized to the MNI space. Once all participants’ p.a._norm_ images were warped to the MNI space, the mean p.a._norm_ image across *N*=8 participants for each of the four DDE sets was computed and used for the CTI fit (hereafter referred to as *MNI-space CTI analysis*). Two subjects were excluded for the group MNI space analysis: i) one because of enlarged ventricles, and ii) because of unsuccessful alignment at the registration process.

Registrations’ warp fields and matrices were also applied to the above listed FreeSurfer-derived ROIs (see *Anatomical images* Section), to warp them to MNI space. For each ROI, the intersection voxels across all *N*=8 included participants were defined as ROIs for the MNI-space CTI analysis (hereafter referred to as ROIs_MNI_), except for GM where voxels were included if common to at least six participants to get a more inclusive mask. Values derived from the MNI-space CTI analysis were then extracted from each ROI_MNI_ for the quantitative analysis of the Kurtosis sources and outlier values were removed with the isoutlier function (method: Grubbs) in MatLab R2017b. Finally, in order to assess the extent of each Kurtosis source contribution to *K*_t_ maps, the ratio between each Kurtosis source and *K*_t_ was computed, and values were extracted from each ROI for further analysis. Voxels with *K* ≤ 0 (any kurtosis source) were excluded from this analysis (the ratio of excluded voxels on total voxels can be found in Fig. S1).

### Correlation Tensor Imaging fit

The CTI fit procedure (described in Henriques et al., 2020a, 2021) was performed using the linear least square fit function in MatLab (version R2017b). The CTI fit generates the following maps: *D*_t_, *K*_t_, *K*_aniso_, *K*_iso_, and *μK* maps. For each of these maps, values were extracted from each FreeSurfer-derived ROI (see *Anatomical images* Section) aligned to the diffusion space. Outlier values were removed with the isoutlier function (method: Grubbs) in MatLab R2017b, and mean and standard deviation (SD) values were computed per each ROI. For the phantom, values within an eroded phantom mask excluding voxels closer than 18 mm to the phantom surface were extracted.

### MNI-space Multiple Gaussian Component (MGC) assumption analysis

To assess the effect of neglecting *μK* on the other Kurtosis sources and thus the validity of the MGC assumption, similarly to Henriques et al., 2021c, *K*_aniso_ and *K*_iso_ were derived under the MGC assumption according to the tensor-valued information of the diffusion MRI acquisitions (Westin et al., 2014; Szczepankiewicz et al., 2019a). p.a._norm_ images used for the *MNI-space CTI analysis* (thus only using axial tensor-valued experiments), were fitted with eq. (5):

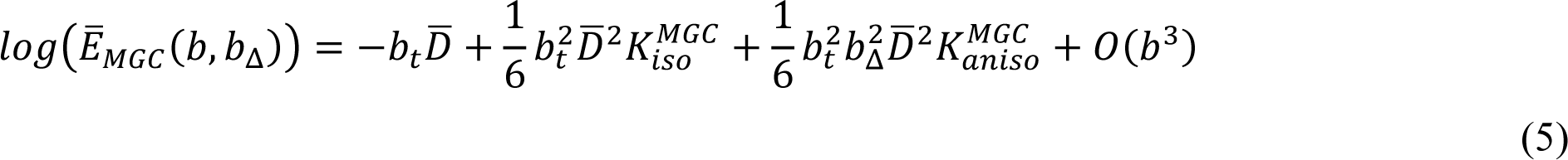

with *b*_t_ being the cumulative *b*-value (*i.e.* considering both pairs of diffusion gradients), and *b*_Δ_ being 1 for DDE sets #1-3 and −1/2 for DDE set #4.

#### MNI-space single-polarity CTI analysis

In order to assess effects of imaging cross-terms, p.a._norm_ was computed separately on data corresponding to each polarity of the diffusion gradients (*i.e.* “positive” and “negative”, Fig. 1B, bottom). p.a._norm_ images then underwent the same registration pipeline described in the *Registration to MNI space and MNI-space CTI analysis* paragraph. *μK* values were extracted from each ROI for their quantitative analysis.

### Data and code availability

Data and code used for data analysis are available via request to the authors, with the need of a formal sharing agreement.

## Results

### Data quality

Fig. 2A shows an example b=0 volume with overlaid FreeSurfer-derived ROIs aligned to the diffusion space for a representative subject. Across subjects, mean SNR ± one SD within the brain was 20 ± 1.35. The SNR map and p.a._norm_ images of non-smoothed data for a representative subject are shown in Fig. 2B and C, respectively. The difference between parallel and antiparallel gradient pairs designed to test for the long mixing time regime, is shown in Fig. 2D: the signal differences did not reveal visible anatomical structures for all sampled directions, and were on the order of the tSD estimated on b=0 volumes in the same series.

### Phantom assessments

CTI maps from the phantom (designed to ensure no systematic effects are observed) are shown in Fig. 2E (before Gaussian smoothing). Mean ± one SD values per each CTI map were: *D*_t_: 2.16 ± 0.11; *K*_t_: 0.09 ± 0.04; *K*_aniso_: 0.00 ± 0.01; *K*_iso_: 0.08 ± 0.04; *μK*: 0.01 ± 0.01; (see also Table 1).

**Table 1.** CTI maps values for individual human and phantom data. Values correspond to mean ± one SD. For human data, the SD was computed across mean values of each participant’s data. For phantom data, the SD was computed spatially, *i.e.* across all included phantom voxels. Abbreviations: CSF: Cerebrospinal fluid in lateral ventricles; WM: White Matter; WMCBM: Cerebellar White Matter; GM: Grey Matter; GMCBM: Cerebellar Grey Matter; AMG: Amygdala; Cd: Caudate; HPC: Hippocampus; GP: Globus Pallidus; PU: Putamen; TH: Thalamus.

| Map | Human data |  |  |  |  |  |  |  |  |  |  | Phantom |
| --- | --- | --- | --- | --- | --- | --- | --- | --- | --- | --- | --- | --- |
|  | Regions of Interest (ROIs) |  |  |  |  |  |  |  |  |  |  |  |
|  | CSF | WM | WM<sub>CBM</sub> | GM | GM<sub>CBM</sub> | AMG | Cd | HPC | GP | PU | TH |  |
| $K_t$ | 0.49 ± 0.04 | 1.04 ± 0.02 | 1.23 ± 0.06 | 0.78 ± 0.01 | 0.97 ± 0.04 | 0.82 ± 0.04 | 0.83 ± 0.06 | 0.86 ± 0.03 | 1.54 ± 0.08 | 1.08 ± 0.05 | 1.1 ± 0.07 | 0.09 ± 0.04 |
| $K_{aniso}$ | 0.02 ± 0.01 | 0.40 ± 0.03 | 0.48 ± 0.03 | 0.08 ± 0.01 | 0.10 ± 0.02 | 0.07 ± 0.01 | 0.06 ± 0.02 | 0.06 ± 0.01 | 0.26 ± 0.08 | 0.15 ± 0.03 | 0.21 ± 0.02 | 0.00 ± 0.01 |
| $K_{iso}$ | 0.46 ± 0.03 | 0.47 ± 0.04 | 0.60 ± 0.07 | 0.57 ± 0.02 | 0.70 ± 0.05 | 0.60 ± 0.04 | 0.65 ± 0.05 | 0.69 ± 0.02 | 1.13 ± 0.13 | 0.73 ± 0.08 | 0.80 ± 0.08 | 0.08 ± 0.04 |
| $\mu K$ | 0.01 ± 0.01 | 0.16 ± 0.01 | 0.15 ± 0.03 | 0.13 ± 0.02 | 0.18 ± 0.02 | 0.16 ± 0.01 | 0.12 ± 0.02 | 0.10 ± 0.01 | 0.15 ± 0.04 | 0.20 ± 0.02 | 0.10 ± 0.02 | 0.01 ± 0.01 |

### Individual CTI maps analysis

All data was successfully acquired without any interruption due to uncooperative participants. Individual CTI-derived maps are shown in Fig. 3. Between subjects, maps consistently showed larger *K*_aniso_ values for white matter areas, larger *K*_iso_ values at the interface between tissues, and *μK* values that were centered on zero for CSF and non-vanishing positive for both grey and white matter. Across-subject mean ± one SD Kurtosis values of CTI-derived maps are listed per each ROI in Table 1. Distributions of Kurtosis values on individual maps per each ROI can be found on Fig. S2.

**Fig. 3.**
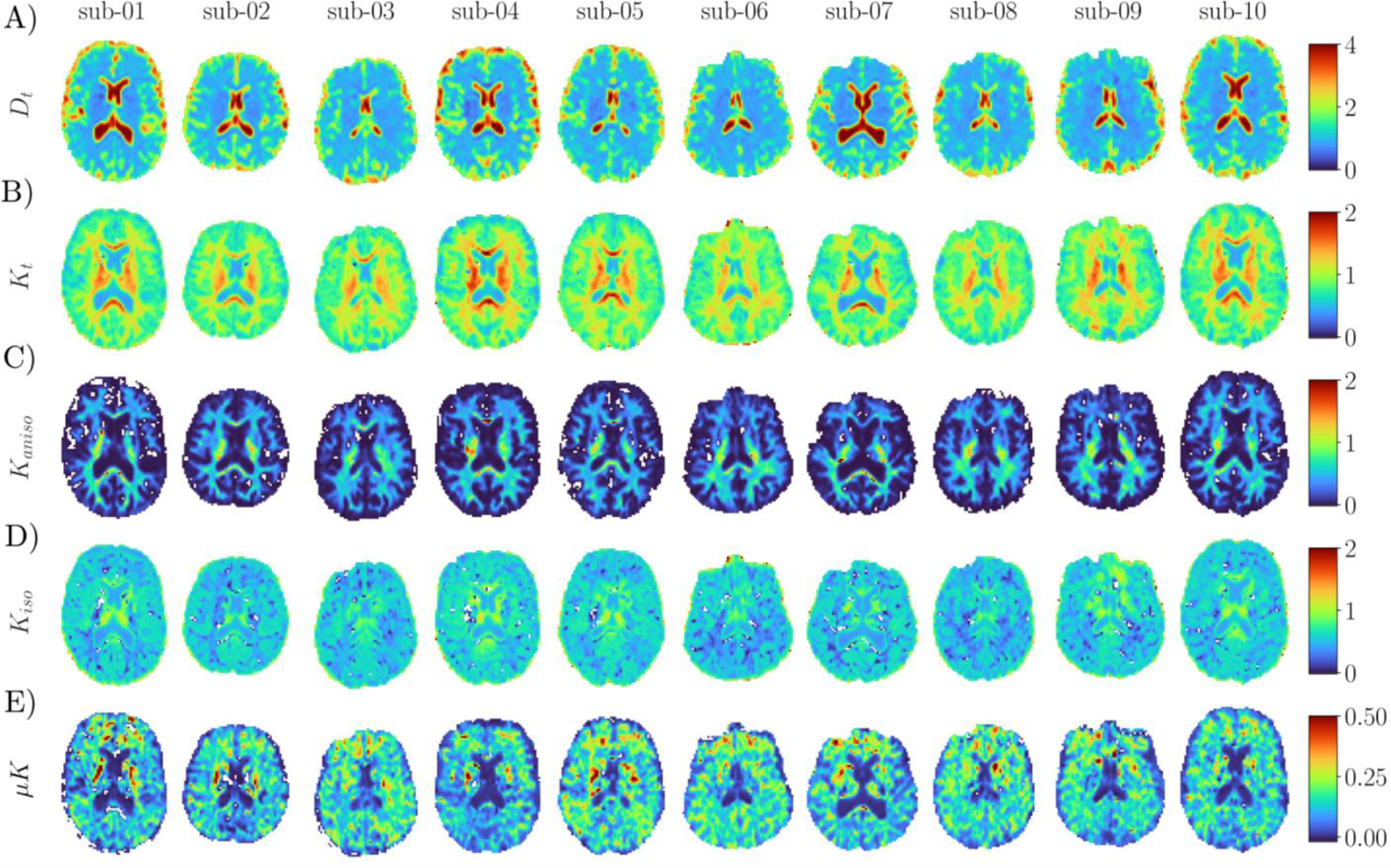
Individual CTI-derived maps in native space. **A)** *D*t maps; **B)** *K*t maps; **C)** *K*aniso maps; **D)** *K*iso maps. White voxels within the brain correspond to negative *K*iso values, possibly associated with noise effects; a discussion on negative kurtosis values can be found *e.g.* in Henriques et al., 2021d; **E)** *μK* maps. Note the different ranges in the colorbars for different CTI metrics.

### MNI-space CTI analysis

CTI maps computed on across-subjects averaged p.a._norm_ images are shown in Fig. 4A, along with distributions of values in each ROI (Fig. 4B). Consistently with individual-level maps, *K*_aniso_ was larger in WM regions, and *K*_iso_ was larger in regions that likely present higher degree of partial volume with CSF (regions near ventricles and edge of cortex); *μK* showed zero-centered values in CSF, but non-vanishing positive values both in grey and white matter (Table 2, voxels identified as outliers and excluded from the plotted distributions and quantitative analyses were <3% for all ROIs and maps except for GP *K*_aniso_, where excluded voxels represented 3.64% of total voxels; see Table S1 for number of excluded voxels and percentages on total voxels for all ROIs and maps). Fig. 5A-C shows maps of the ratio between each Kurtosis source and *K*_t_, along with barplots showing the Kurtosis components per each ROI (Fig. 5D): *K*_iso_ appears to be the largest Kurtosis source for all tissues except white matter regions, with *μK* accounting for 8-20% of *K*_t_ in all ROIs except for CSF.

**Fig. 4.**
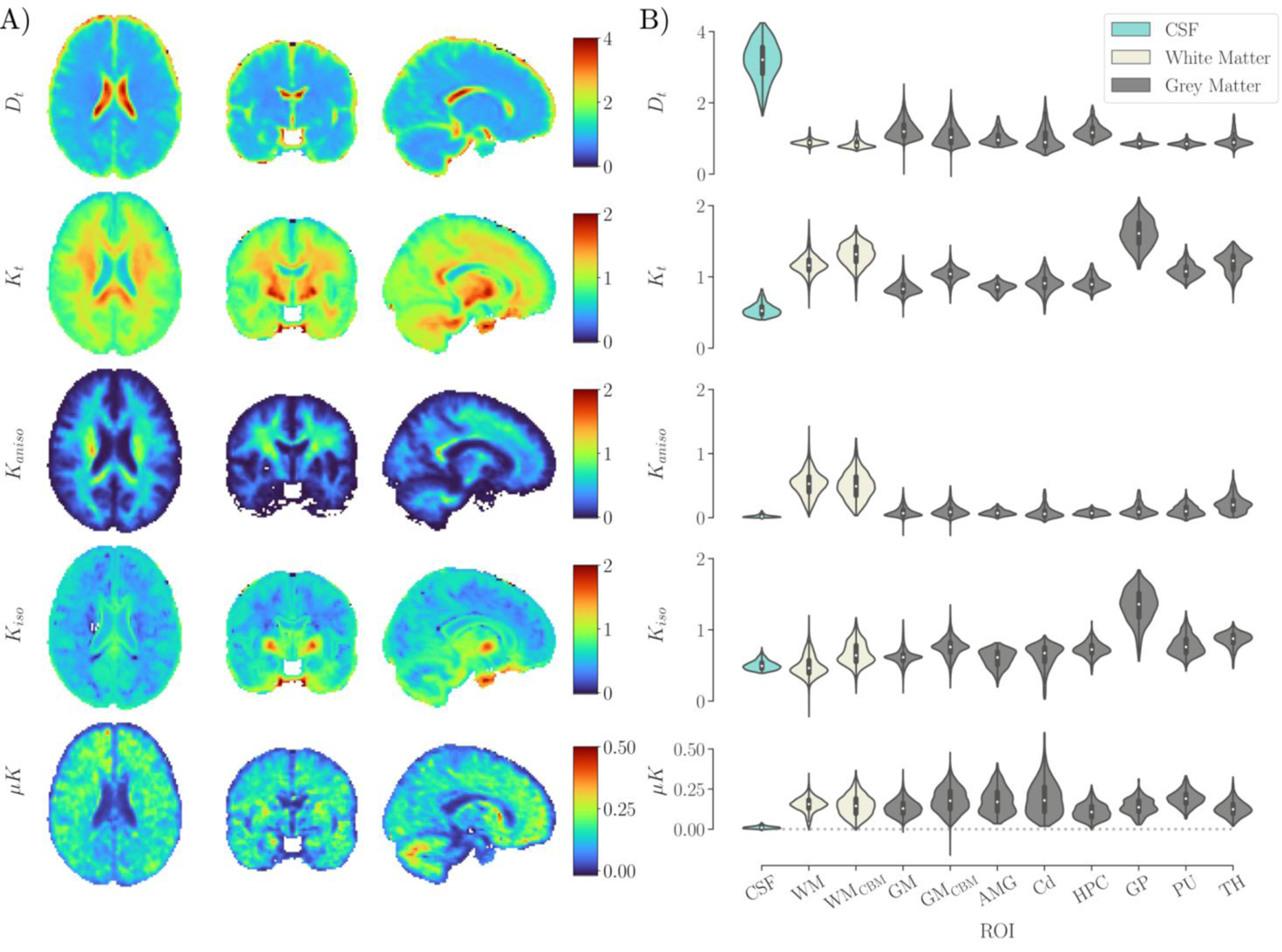
, CTI maps derived from the average of data from *N*=8 subjects in MNI space (MNI-space CTI analysis). **A)** Axial, coronal, and lateral views of *D*t, *K*t, *K*aniso, *K*iso, and *μK* maps derived by fitting the averaged data of *N*=8 subjects, previously normalized to the 2mm MNI atlas. Note the different ranges in the colorbars for different CTI metrics. **B)** Distributions of values per each CTI metric in each Region of Interest (ROI) considered in this study. CSF: Cerebrospinal fluid in lateral ventricles; WM: White Matter; WMCBM: Cerebellar White Matter; GM: Grey Matter; GMCBM: Cerebellar Grey Matter; AMG: Amygdala; Cd: Caudate; HPC: Hippocampus; GP: Globus Pallidus; PU: Putamen; TH: Thalamus.

**Fig. 5.**
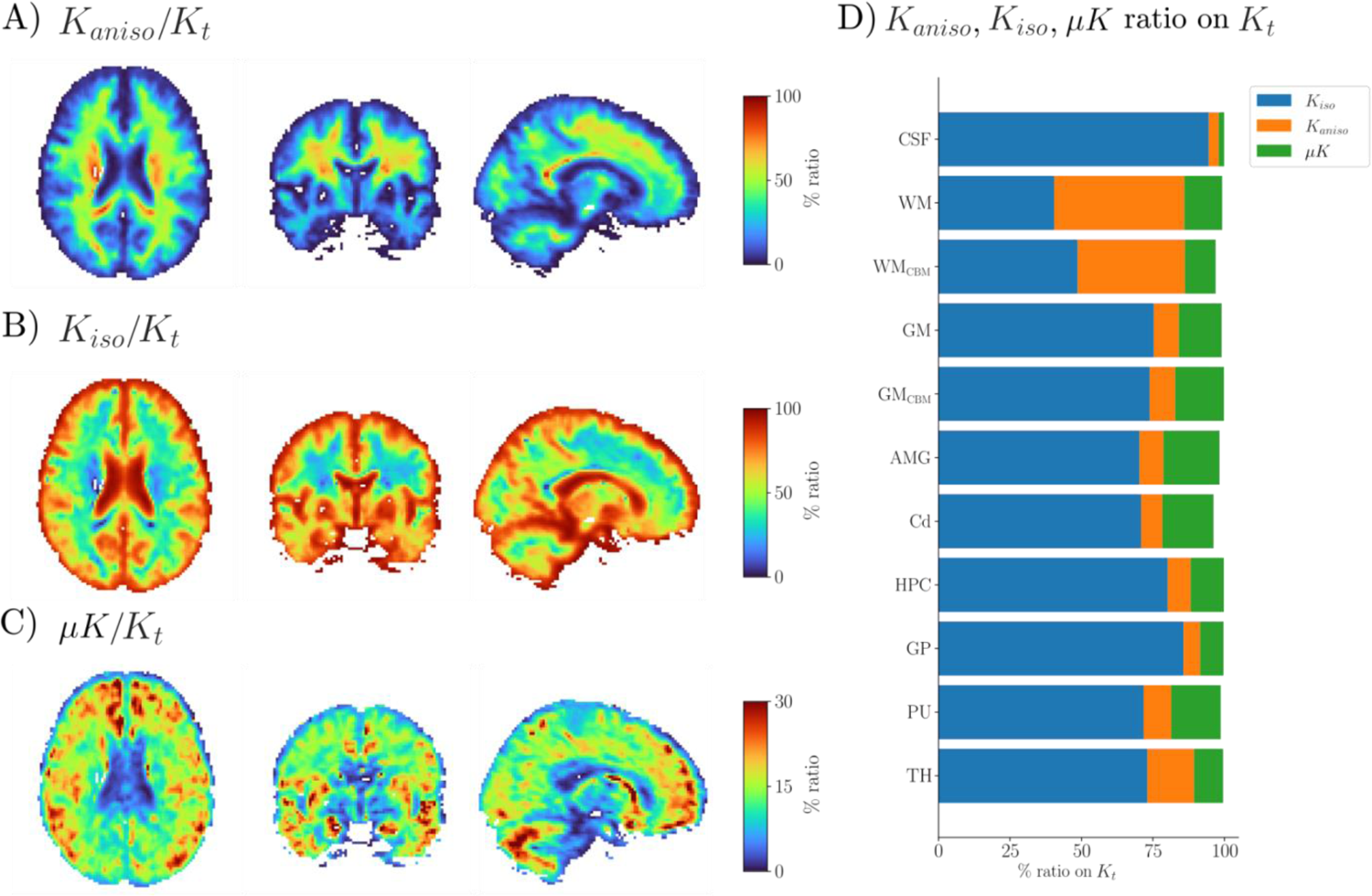
Ratio maps of each Kurtosis source on total Kurtosis. CTI maps derived from the average of data from *N*=8 subjects in MNI space. Axial, coronal, and lateral views of **A)** *K*aniso/*K*t ratio, **B)** *K*iso/*K*t ratio, and **C)** *μK*/*K*t ratio. Note the different ranges in the colorbars for different ratios. **D)** Barplot showing median percent ratio per each Kurtosis source per each Region of Interest (ROI) considered in this study. CSF: Cerebrospinal fluid in lateral ventricles; WM: White Matter; WMCBM: Cerebellar White Matter; GM: Grey Matter; GMCBM: Cerebellar Grey Matter; AMG: Amygdala; Cd: Caudate; HPC: Hippocampus; GP: Globus Pallidus; PU: Putamen; TH: Thalamus.

**Table 2.** Kurtosis sources values for CTI maps derived from the average of data from *N*=8 subjects in MNI space (MNI-space CTI analysis). Values of Kurtosis sources (*K*aniso, *K*iso, *μK*) correspond to mean ± one SD. Values of percent ratio between each Kurtosis source and *K*t correspond to median (interquartile range, IQR). The SDs and the IQRs were computed spatially, *i.e.* across all included ROI voxels. Abbreviations: CSF: Cerebrospinal fluid in lateral ventricles; WM: White Matter; WMCBM: Cerebellar White Matter; GM: Grey Matter; GMCBM: Cerebellar Grey Matter; AMG: Amygdala; Cd: Caudate; HPC: Hippocampus; GP: Globus Pallidus; PU: Putamen; TH: Thalamus.

| Map | Human group average template data: *MNI-space CTI analysis* |  |  |  |  |  |  |  |  |  |  |
| --- | --- | --- | --- | --- | --- | --- | --- | --- | --- | --- | --- |
|  | Regions of Interest (ROIs) |  |  |  |  |  |  |  |  |  |  |
|  | CSF | WM | WM<sub>CBM</sub> | GM | GM<sub>CBM</sub> | AMG | Cd | HPC | GP | PU | TH |
| $K_t$ | 0.54 ± 0.09 | 1.17 ± 0.13 | 1.31 ± 0.15 | 0.84 ± 0.09 | 1.04 ± 0.09 | 0.85 ± 0.07 | 0.91 ± 0.12 | 0.91 ± 0.09 | 1.61 ± 0.19 | 1.09 ± 0.1 | 1.20 ± 0.14 |
| $K_{\text{aniso}}$ | 0.02 ± 0.02 | 0.52 ± 0.19 | 0.50 ± 0.21 | 0.90 ± 0.08 | 0.10 ± 0.09 | 0.08 ± 0.04 | 0.08 ± 0.09 | 0.07 ± 0.04 | 0.12 ± 0.09 | 0.11 ± 0.08 | 0.21 ± 0.13 |
| $K_{\text{aniso}} / K_t$<br>% ratio | 3.5<br>(3.9) | 45.5<br>(17.2) | 37.5<br>(18.6) | 8.7<br>(9.6) | 9.0<br>(8.7) | 8.3<br>(6.9) | 7.5<br>(9.9) | 8.0<br>(5.8) | 5.8<br>(6.5) | 9.7<br>(11.6) | 16.4<br>(11.3) |
| $K_{\text{iso}}$ | 0.50 ± 0.06 | 0.49 ± 0.15 | 0.67 ± 0.16 | 0.62 ± 0.11 | 0.75 ± 0.13 | 0.60 ± 0.12 | 0.63 ± 0.16 | 0.73 ± 0.1 | 1.34 ± 0.23 | 0.78 ± 0.14 | 0.86 ± 0.1 |
| $K_{\text{iso}} / K_t$<br>% ratio | 94.5<br>(4.9) | 40.4<br>(17.5) | 48.7<br>(20.1) | 75.3<br>(15.5) | 73.8<br>(18.1) | 70.3<br>(18) | 70.8<br>(23.1) | 80.2<br>(10.7) | 85.7<br>(6.5) | 71.7<br>(12.8) | 73.0<br>(11.6) |
| $\mu K$ | 0.01 ± 0.01 | 0.15 ± 0.05 | 0.15 ± 0.07 | 0.13 ± 0.05 | 0.18 ± 0.08 | 0.18 ± 0.08 | 0.19 ± 0.1 | 0.11 ± 0.05 | 0.14 ± 0.05 | 0.19 ± 0.05 | 0.13 ± 0.05 |
| $\mu K / K_t$<br>% ratio | 1.9<br>(1.6) | 13.2<br>(5.1) | 10.8<br>(7.5) | 15.0<br>(7.5) | 17.0<br>(10.1) | 19.6<br>(15.9) | 17.9<br>(16.5) | 11.6<br>(7.5) | 8.2<br>(4) | 17.3<br>(6.4) | 10.1<br>(5.1) |

#### MNI-space CTI vs. MGC analysis

Fig. 6 shows the relationship between total, anisotropic and isotropic kurtosis estimates derived from CTI and the corresponding ones estimated under the MGC assumption, which neglects *μK*. For any of these metrics, a bias between the two approaches will be seen as a distribution that departs from the diagonal. Relative to CTI, the MGC approach leads to lower *K*_t_ in voxels with high *μK*. The opposite pattern was observed for *K*_aniso_ and *K*_iso_, where estimates derived under the MGC assumption were overestimated compared to CTI-derived values, with larger deviations being associated to larger *μK* values.

**Fig. 6.**
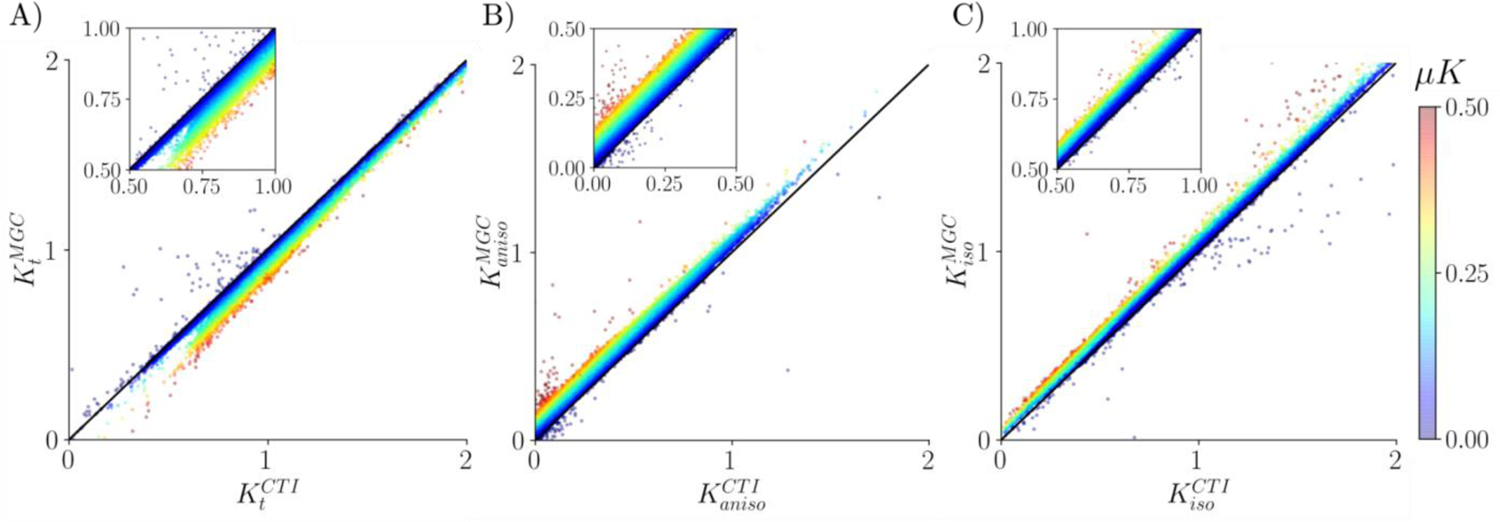
Effect of neglecting *μK* on *K*t, *K*aniso, and *K*iso. CTI- and MGC-derived maps were estimated on the average of data from *N*=8 subjects in MNI space. **A)** CTI-vs. MGC-derived *K*t. **B)** CTI-vs. MGC-derived *K*aniso. **C)** CTI-vs. MGC-derived *K*iso. Points are color-coded according to their *μK* value, inset plots are added for better visibility.

### MNI-space single-polarity CTI analysis

Fig. 7 shows CTI-derived maps from the averaged data corresponding to either both or single repetitions of the CTI acquisition. In comparison with maps estimated from two repetitions (corresponding to a ∼52 minutes-long acquisition), maps computed from single repetitions suggest that there is no systematic bias when considering only one polarity of the diffusion gradients. Distributions of *μK* values derived from single repetitions per each ROI can be found on Fig. S3.

**Fig. 7.**
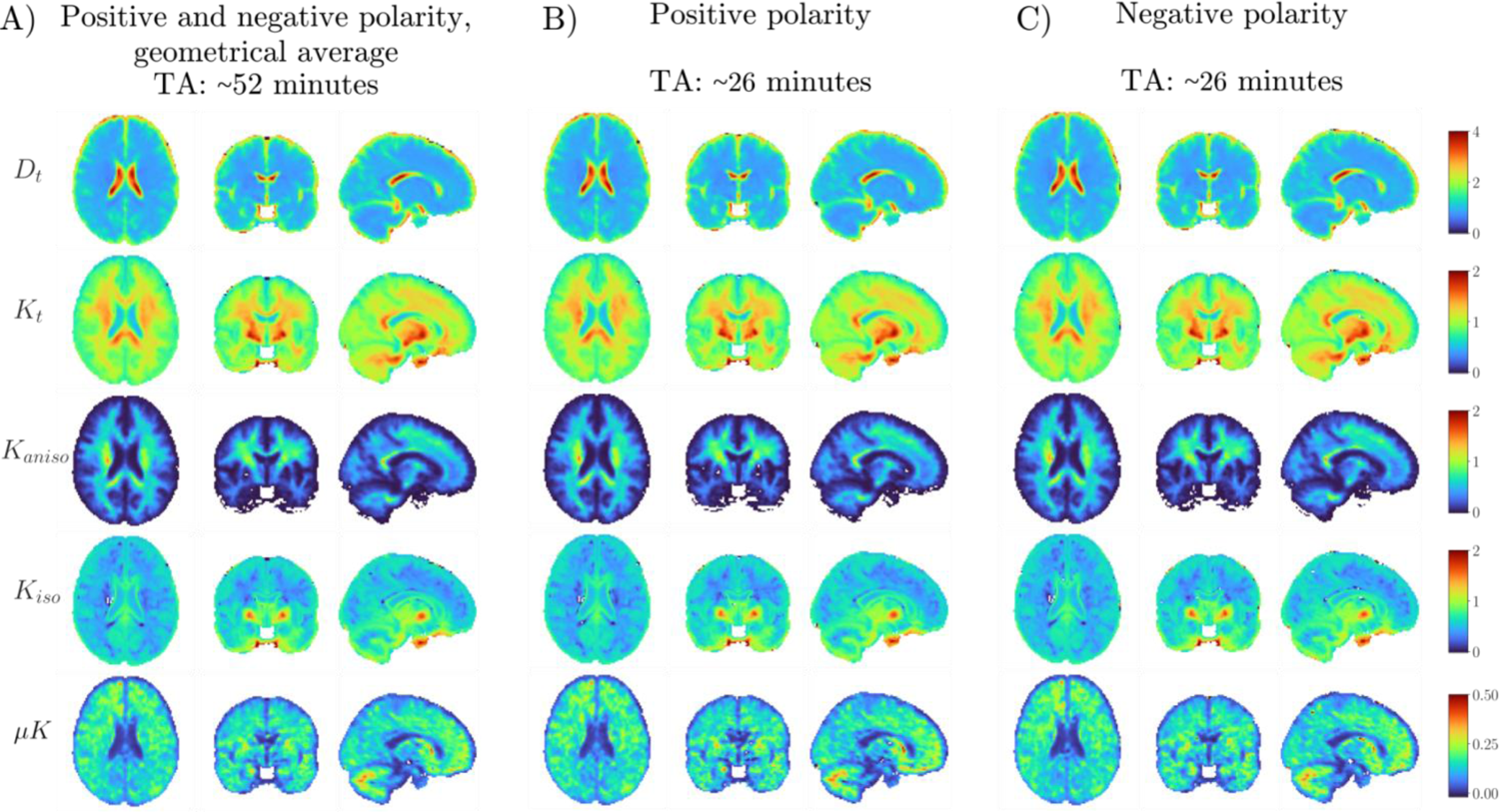
Effect of reducing the acquisition time by 50% on CTI-derived maps. CTI-derived maps were estimated on the average of data from *N*=8 subjects in MNI space. Axial, coronal, and lateral views of **A)** CTI-derived maps estimated from the geometrical average of two repetitions, each acquired with opposite polarity of the diffusion gradients (here referred to as “positive” and “negative”, Neeman et al., 1991; Ianuş et al., 2018), yielding a 52-minutes long acquisition; **B)** CTI-derived maps estimated from only one repetition corresponding to the “positive” polarity of the diffusion gradients; **C)** CTI-derived maps estimated from only one repetition corresponding to the “negative” polarity of the diffusion gradients. Note the different ranges in the colorbars for different CTI metrics. Acquisition time corresponding to only one repetition (panels B and C) was 26 minutes. TA: Acquisition time.

## Discussion

Disentangling kurtosis sources in biological systems is attracting increasing interest given the potential to increase specificity and potentially provide novel non-invasive quantitative markers of microstructural properties *in vivo*. The recently-proposed CTI framework is designed to resolve anisotropic, isotropic, and microscopic kurtosis sources, which indeed provided much insight in preclinical imaging. In this work, we have extended the CTI methodology towards human imaging on a clinical 3T MRI system and aimed to investigate the kurtosis sources in healthy volunteers. Our main findings revealed not only the anisotropic and isotropic kurtosis sources in the human brain, but also clearly showed non-vanishing positive microscopic kurtosis contributions accounting for 8-20% of the total diffusional kurtosis, larger in white matter than in gray matter, and vanishing in the ventricles. Consistently with this, we also show that MGC-based approaches for kurtosis source separation are biased when neglecting microscopic kurtosis. The results of the present study pave the way towards quantifying kurtosis sources more precisely and accurately, investigating them in different types of disease. Our findings motivate further developments to accelerate data acquisition to make human brain CTI more compatible with clinical scan times.

### Value of separating kurtosis sources using the CTI methodology

Decomposing the total kurtosis into its underlying sources using CTI revealed specific complementary contrasts for each kurtosis component both in individual maps and the group average template. For instance, *K*_t_ maps show elevated values in regions where partial volume effects are expected, such as around the lateral ventricles, and more in general at tissue interfaces; the separation in sources, however, allows to disentangle the relative contributions arising from each kurtosis source. In particular, the CTI method enables the evaluation of *μK* which cannot be obtained from other tensor-valued approaches. More specifically, with the diffusion times adopted in the current study, positive *μK* values were observed both for grey and white matter for both individual maps and the group average template, suggesting that the kurtosis component arising from non-Gaussian effects might be ubiquitous in the cerebral tissue. Interestingly, the quantitative analysis of both individual maps and the group average template also showed large *K*_t_ values for the globus pallidus, and the separation of *K*_t_ in its sources revealed that such elevated kurtosis values were mostly driven by high *K*_iso_. The short *T*_2_ characterizing this structure in combination with the longer echo time needed for the DDE preparation, yields a low SNR, which is likely the root of the observed high *K*_iso_ values (Jensen & Helpern, 2010). Nevertheless, the observations of expected contrasts, together with the overall agreement between our average white and grey matter *μK* estimates and those obtained by Dhital et al., 2018, strongly support the feasibility of resolving the kurtosis sources according to the CTI methodology in clinical systems.

Disentangling diffusional kurtosis sources without *a priori* assumptions on the diffusion mode in the tissue represents a new advancement from two different perspectives. On the one hand, it represents an opportunity to enhance specificity in microstructural imaging: the conflation of mesoscopic and microscopic effects in conventional DKI metrics renders the method highly unspecific; separating the sources using CTI thus provides a way to bypass this limitation and quantify the contribution of different sources to the total kurtosis. This could facilitate the detection of different biological processes that may not be visible in conventional DKI due to their counterbalancing effects on overall tissue heterogeneity (*e.g.* decreased fiber dispersion and fiber anisotropy due to fiber degradation leading to decreased *K*_aniso_, and concomitant variations in the extracellular matrix or the intracellular space due to vasogenic or cytotoxic phenomena leading to increased *K*_iso_ and variations in *μK*). Thus, the possibility to separately map each kurtosis source could reveal the relative “weights” contributing to overall tissue heterogeneity. In turn, this may enhance the biological interpretation of the underlying pathophysiology, as already observed for predicting tumor histology (Szczepankiewicz et al., 2015; Szczepankiewicz et al., 2016; Nilsson et al., 2020) and investigating the mechanisms of the early tissue responses to ischemia (Alves et al., 2021). On the other hand, from the perspective of the biophysical modeling of the dMRI signal, signal representations could play a crucial role for the development of dMRI microstructural models: by providing an unbiased picture of signal features, indeed signal representations may help to set the ground for establishing adequate priors to then be adopted in the modeling process, with the aim of further determining valid links between the detected signal features and their biological underpinnings (Novikov et al., 2018). Moreover, the increasingly available “orthogonal” diffusion-based contrasts, such as MDE preparations, together with the exploration of additional dimensions as the diffusion time, have indeed already helped to identify biases, suggest priors, and constructively shape the ranges of validity of different representations and models (*e.g.* De Santis et al., 2016; Lampinen et al, 2017; Jespersen et al., 2018; Henriques et al., 2019; Lampinen et al., 2019; Henriques et al., 2021c), thus yielding insightful information in the progress toward the validation and development of such approaches (Alexander et al., 2017; Dyrby et al., 2018; Jespersen, 2018).

### Comparison of CTI and MGC-driven approaches

Several studies have highlighted the limitations and the ambiguity of the SDE PGSE signal in resolving the kurtosis sources (*e.g.* Henriques et al., 2019), and until very recently MDE approaches relying on the MGC assumption represented the only option to separate the kurtosis sources, implicitly assuming *μK* = 0. Using the CTI methodology, our study shows clear evidence for residual non-Gaussian diffusion in the healthy adult human brain (the *μK* component, previously referred to as intra-compartmental kurtosis in Henriques et al., 2020), both at the individual level and at the MNI space analysis. The vanishing of *μK* in CSF lends further credence to our measurements, and suggests it cannot be ignored in the human brain, even when using a 3 T clinical MRI setting. In the brain tissue, the quantitative analysis of the relative weight of the *μK* component on the total kurtosis (in the MNI space analysis) suggests that *μK* accounts for between 8 and 20% of total kurtosis. In other words, at the length scales probed by the diffusion times achievable in clinical systems (in our system *G*_max_ = 80 mT/m), non-Gaussian diffusion components might represent up to 20% of the total Kurtosis. Systems with higher performance gradients, capable to achieve similar diffusion weightings in shorter diffusion time, might potentially reveal even larger weights of the *μK* component, as observed e.g. in Henriques et al., 2021c, in the preclinical setting. Nevertheless, in the current work by assessing the impact of neglecting *μK* in the estimation of *K*_aniso_ and *K*_iso_, *i.e.* under the MGC assumption, consistently with the findings in the rat brains the preclinical setting in Henriques et al., 2021c, our data show lower estimates for *K*_t_ and larger estimates for *K*_aniso_ and *K*_iso_ under the MGC assumption, with larger biases for larger (neglected) *μK* values. As highlighted by the results in Jespersen et al., 2019, it is imperative to carefully assess the effects of non-Gaussian diffusion on the MGC assumption. In addition, the *μK* may become much more dominant upon disease (c.f. Alves et al., 2021). We believe that the results presented in the current work are crucial for suggesting an initial range of potential bias associated with the MGC assumption in the healthy human brain at the diffusion times adopted in this study, and thus to help the interpretation of *K*_aniso_ and *K*_iso_ estimates derived under frameworks assuming Gaussian diffusion over different tissue domains.

### μK as a new source of contrast

Importantly, mapping the *μK* component, in addition to informing the frameworks neglecting non-Gaussian diffusion, represents a new source of contrast *per se*. By definition, indeed, the *μK* component represents the residual non-Gaussianity not captured by *K*_aniso_ and *K*_iso_, which our current results suggest to be ubiquitous in the human cerebral tissue. In addition to studies directly investigating *μK*, insights on residual non-Gaussian diffusion mainly come from the works investigating diffusion and in turn kurtosis time-dependence. Differently from the CTI approach, these require multiple measurements at different diffusion times, which can be onerous for clinical scanning. The observation of time-dependent diffusion, and in turn of kurtosis, coefficients indeed suggests that at least one tissue compartment exhibits non-Gaussian diffusion (Fieremans et al., 2016; Novikov et al., 2019; Lee et al., 2020a). Such deviations from Gaussianity are believed to be associated with structural disorder and cross-sectional variance characterizing the sampled tissue (Novikov et al., 2014, see also below), and to be modulated by exchange effects (Nilsson et al., 2013 - both intra-cellular and intra-extracellular exchange, with the latter potentially dominating at the diffusion times adopted in the current studies, Ianuş et al., 2021). It is important to point out that since the length scale of the displacements sensed by the diffusion encoding depends on the diffusion time (Δ) adopted (in addition to the substrate diffusion coefficient), at very long diffusion times the *μK* component might vanish due to the substrate’s complete coarse-graining. In this situation, diffusion in each non-exchanging compartment then averages out and may be described as a Gaussian, and thus by a uniform effective diffusion coefficient (for a description of the diffusion phenomenon as a coarse-graining process see Novikov et al., 2019). Conversely, the *μK* component is expected to be increasingly important at short diffusion times, a regime at which more pronounced time-dependency has been observed (*e.g.* Fieremans et al., 2016; Jespersen et al., 2018). This short diffusion time regime is more easily achieved by high-performance gradients, such as those available in preclinical and dedicated human MRI systems (McNab et al., 2013b; Jones et al., 2018; Fan et al., 2020; Lee et al., 2020b; Henriques et al., 2021c; Huang et al., 2021).

Several pieces of evidence both from preclinical and clinical applications suggest the diagnostic potential of getting insights in such non-Gaussian diffusion arising from restrictions or structural disorder. For instance, in the preclinical setting, mapping the *μK* component in the rat brain post-ischemia revealed enhanced sensitivity to stroke regions and allowed more specific insights into cellular mechanisms involved in response to stroke (Alves et al., 2021).

In clinical systems, insights into such non-Gaussian diffusion in the human brain so far mainly come from studies investigating diffusion and kurtosis time-dependency. While several studies did not observe time-dependent diffusion effects in the *in vivo* human brain (Clark et al., 2001; Nilsson et al., 2009), other studies reported diffusion time-dependency both for white matter (Horsfield et al., 1994; Baron and Beaulieu, 2014; Van et al., 2014; Fieremans et al., 2016; Lee et al., 2018; Lee et al., 2020b; Arbabi et al., 2020) and grey matter (Baron and Beaulieu, 2014; Lee et al., 2020a). Transverse to axonal bundles, time-dependent diffusion has been observed to report on two-dimensional structural disorder, in turn associated to the extra-axonal space packing geometry (Burcaw et al., 2015; Fieremans et al., 2016; Lee et al., 2018) and to be influenced by variations in axonal caliber, also referred to as axonal beading or varicosities (Ginsburger et al., 2018). Along axonal fibers, a stronger diffusion time- (or frequency-, for OGSE sequences) dependency has been observed to follow the power-law proposed in Novikov et al., 2014 for short-range disorder (Fieremans et al., 2016; Jespersen et al., 2018; Arbabi et al., 2020; Lee et al., 2020b). A link between the observed time-dependent signal modulations and their biological underpinnings comes from simulations of diffusion in three-dimensional reconstructions of histology-derived axonal segments, which clarified that the observed power-law time-dependency along axons arises in association with axonal varicosities (Lee et al., 2020b). Observations for a one-dimensional structural disorder power-law time dependency in grey matter, furthermore, suggest that this might be a universal property of the neural tissue (Does et al., 2003; Novikov et al., 2014; Lee et al., 2020a; Lee et al., 2020b).

### Initial steps toward clinical translation

Data acquisition for this work required almost one hour per participant: two repetitions of the DDE sets required by the CTI methodology were collected, which combined allowed a reduction of effects resulting from cross-terms with imaging gradients. However, in the group template kurtosis maps we found that the distributions of values derived from opposite- or single-polarities were not qualitatively different. This suggests that imaging cross-terms effects may be weak, thus allowing to reduce the acquisition time by half. The resulting ∼30 minutes long acquisition with a single polarity would still be impractical for routine clinical applications, but further work could decrease the number of directions per DDE set to ultimately reduce the scan time further. Nevertheless, we stress that, as pointed out in Henriques et al., 2021c, mapping the *μK* component does not require the acquisition of all the DDE sets required by the CTI methodology: a “direct” estimation of the *μK* component indeed can be accessed via eq. 3, which requires the acquisition of just two DDE sets out of the four required by the complete CTI methodology, together with data for estimating *D*_t_. A single repetition of such an acquisition in our protocol would require ∼15 minutes, which despite being a long scan time for some clinical applications, is closer to an acceptable acquisition time. Anyways, future work should further address the test-retest reliability, suggest normative values at specific diffusion times on a larger sample, and investigate the sensitivity and specificity of the *μK* component in physiological (*e.g.* aging) and disease processes.

### Limitations

This work has some limitations. For our phantom validation, we adopted an isotropic homogeneous phantom characterized by Gaussian diffusion. While our results confirm the expected Gaussian-only diffusion both in the phantom and in the lateral ventricles in the brain, the development of realistic phantoms with different levels of disordered structures mimicking the salient microstructural features believed to be associated with the *μK* component would be beneficial and support the current findings. The development of phantoms is an active and crucial branch in the field of the study of microstructure with dMRI (*e.g.* Shemesh et al., 2010; Shemesh et al., 2012a; Nilsson et al., 2017; Fieremans and Lee, 2018; Giménez et al., 2018). Several studies have used numerical phantoms to investigate dMRI signal modulations associated with non-Gaussianity arising from restrictions or structural disorder (Ginsburger et al., 2018; Palombo et al., 2018; Henriques et al., 2020; Lee et al., 2020b; Lee et al., 2020c; Henriques et al., 2021c; Alves et al., 2021). However, increasing evidence for the accessibility to these microscopic disorder features via dMRI should prompt new crucial advancements in the manufacturing of physical phantoms allowing a further validation of the multiple kurtosis sources.

In addition, when performing the demonstrative MNI-space single-polarity CTI analysis, data were preprocessed considering both polarities. Future studies should investigate further the impact of reducing the number of directions on the CTI metrics estimation by considering accordingly the subset of data in the whole preprocessing stream.

Another limitation is that the current framework assumes negligible exchange between tissue microdomains. More specifically, it should be noted that the microscopic kurtosis is currently estimated as the subtraction of the kurtosis arising from isotropic and anisotropic variances from the total kurtosis, leaving thus the possibility of exchange as a potential contributor. In other words, when two components (even if gaussian) are exchanging in the time-scale of the diffusion experiment (diffusion time, mixing time in the case of CTI), then the total kurtosis and the Z-tensor will exhibit different terms associated with the exchange rates. On subtraction, the difference between these terms can generate a finite exchange-driven microscopic kurtosis. Thus, in principle, the microscopic kurtosis contrast could reflect kurtosis arising from cross sectional variance (positive *μK*), exchange, or a combination of both. Future studies should thus be designed to further disentangle these two effects and investigate more deeply the biological underpinnings of *μK* in the human brain. Nevertheless, separating the kurtosis arising from microscopic sources and variance in tensor magnitude and anisotropy is expected to be highly useful even before all the specific underpinnings are fully resolved (c.f. the sensitivity of microscopic kurtosis contrast in stroke).

## Conclusion

This work demonstrates the translation of the CTI methodology from the preclinical to the clinical MRI setting, prompted by the increasing evidence suggesting the relevance of non-Gaussian diffusion in the characterization of the human brain microstructure. While non-Gaussian effects have been typically investigated by varying diffusion times, making such acquisitions potentially long and impractical, the CTI methodology offers a more practical alternative which has the potential to be further optimized for clinical applications. Mapping the microscopic kurtosis in human brain tissue for the first time revealed that, while until now commonly neglected, this component is non-vanishing. Consistent with this, we show that ignoring microscopic kurtosis affects the estimation of the other kurtosis components. The possibility of mapping microscopic kurtosis in humans opens an intriguing new window on microscopic tissue features of great clinical and neuroscientific interest.

## Acknowledgements

The authors thank Stefano Tambalo for support with data acquisition. This research was supported by the Caritro Foundation, Italy, the European Research Council (ERC) under the European Union’s Horizon 2020 research and innovation programme (Starting Grant, agreement No. 679058), “la Caixa” Foundation (ID 100010434), and European Union’s Horizon 2020 research and innovation programme under the Marie Skłodowska-Curie grant agreement No 847648, fellowship code CF/BQ/PI20/11760029.

