## Supplementary Materials for "In vivo Correlation Tensor MRI reveals microscopic kurtosis in the human brain on a clinical 3T scanner"

### Table S1

| Map | Human group average template data: *MNI-space CTI analysis* Number and % of voxels identified as outliers on total voxels Regions of Interest (ROIs) | | | | | | | | | | |
|---|---|---|---|---|---|---|---|---|---|---|---|
| | CSF | WM | $WM_{CBM}$ | GM | $GM_{CBM}$ | AMG | Cd | HPC | GP | PU | TH |
| $\boldsymbol{D_t}$ | | | | | | | | | | | |
| count | 0 | 968 | 72 | 22 | 8 | 3 | 1 | 6 | 11 | 15 | 18 |
| % | 0 | 2.78 | 2.79 | 0.05 | 0.07 | 1.02 | 0.14 | 0.86 | 2.67 | 1.37 | 0.95 |
| $\boldsymbol{K_t}$ | | | | | | | | | | | |
| count | 2 | 4 | 3 | 391 | 28 | 1 | 6 | 3 | 0 | 1 | 0 |
| % | 0.21 | 0.01 | 0.12 | 0.95 | 0.26 | 0.34 | 0.82 | 0.43 | 0 | 0.09 | 0 |
| $\boldsymbol{K_{aniso}}$ | | | | | | | | | | | |
| count | 8 | 7 | 0 | 68 | 68 | 1 | 11 | 0 | 15 | 0 | 5 |
| % | 0.82 | 0.02 | 0 | 0.17 | 0.63 | 0.34 | 1.5 | 0 | 3.64 | 0 | 0.26 |
| $\boldsymbol{K_{iso}}$ | | | | | | | | | | | |
| count | 1 | 71 | 0 | 404 | 20 | 0 | 8 | 1 | 0 | 0 | 20 |
| % | 0.1 | 0.2 | 0 | 0.99 | 0.19 | 0 | 1.09 | 0.14 | 0 | 0 | 1.05 |
| $\boldsymbol{\mu K}$ | | | | | | | | | | | |
| count | 2 | 1 | 0 | 33 | 29 | 0 | 0 | 0 | 0 | 0 | 2 |
| % | 0.21 | 0.00 | 0.00 | 0.08 | 0.27 | 0 | 0 | 0 | 0 | 0 | 0.11 |

**Table S1, Count of voxels and percentage on total voxels identified as outliers on maps generated at the MNI-space CTI analysis for the generation of the human group average template maps.** Values identified as outliers were excluded from the subsequent quantitative analyses (plots of distributions of values and mean and SD calculation). Abbreviations: CSF: Cerebrospinal fluid in lateral ventricles; WM: White Matter; $WM_{CBM}$: Cerebellar White Matter; GM: Grey Matter; $GM_{CBM}$: Cerebellar Grey Matter; AMG: Amygdala; Cd: Caudate; HPC: Hippocampus; GP: Globus Pallidus; PU: Putamen; TH: Thalamus.

**Fig. S1**

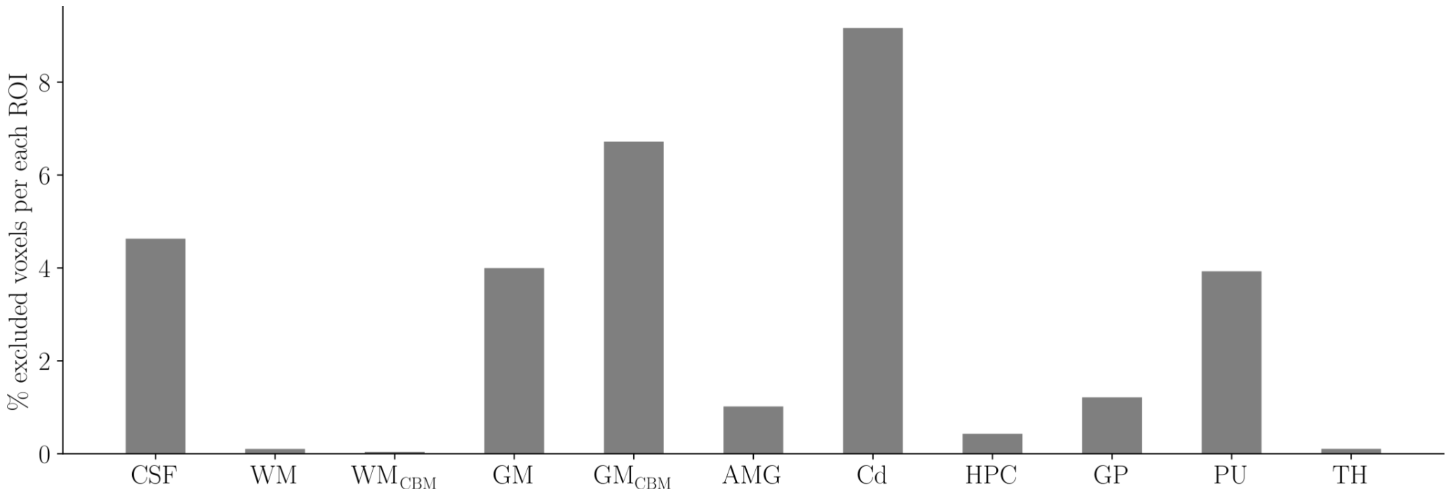


**Fig. S1, Percentage of voxels excluded in the calculation of the % ratio between each kurtosis source and $K_t$.** Voxels were excluded if presenting with $K_t \leq 0$ or negative values in any of the kurtosis sources. Abbreviations: CSF: Cerebrospinal fluid in lateral ventricles; WM: White Matter; $WM_{CBM}$: Cerebellar White Matter; GM: Grey Matter; $GM_{CBM}$: Cerebellar Grey Matter; AMG: Amygdala; Cd: Caudate; HPC: Hippocampus; GP: Globus Pallidus; PU: Putamen; TH: Thalamus.

**Fig. S2**

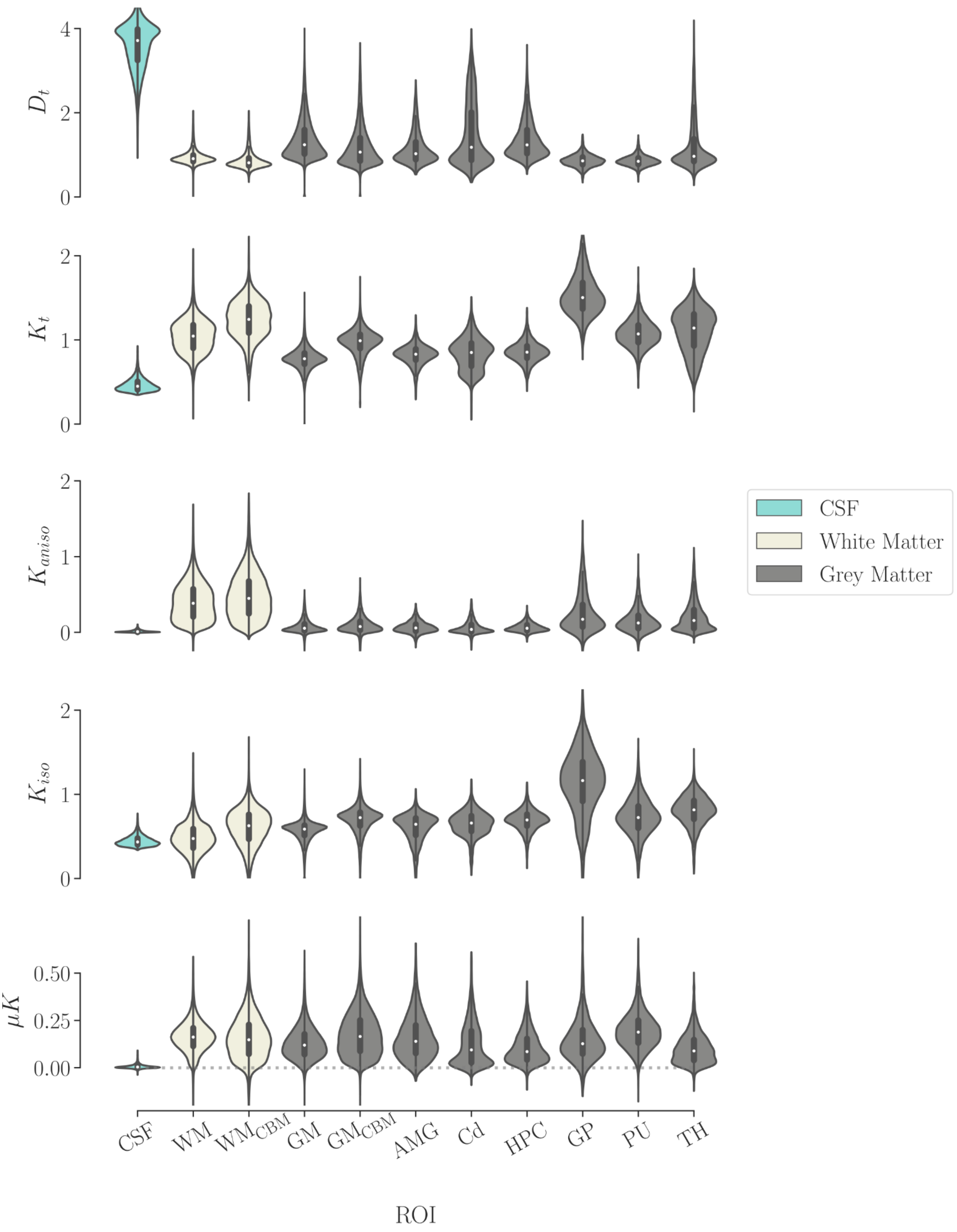


**Fig. S2, Distributions of values from individual CTI maps for each ROI.** Abbreviations: CSF: Cerebrospinal fluid in lateral ventricles; WM: White Matter; $WM_{CBM}$: Cerebellar White Matter; GM: Grey Matter; $GM_{CBM}$: Cerebellar Grey Matter; AMG: Amygdala; Cd: Caudate; HPC: Hippocampus; GP: Globus Pallidus; PU: Putamen; TH: Thalamus.

**Fig. S3**

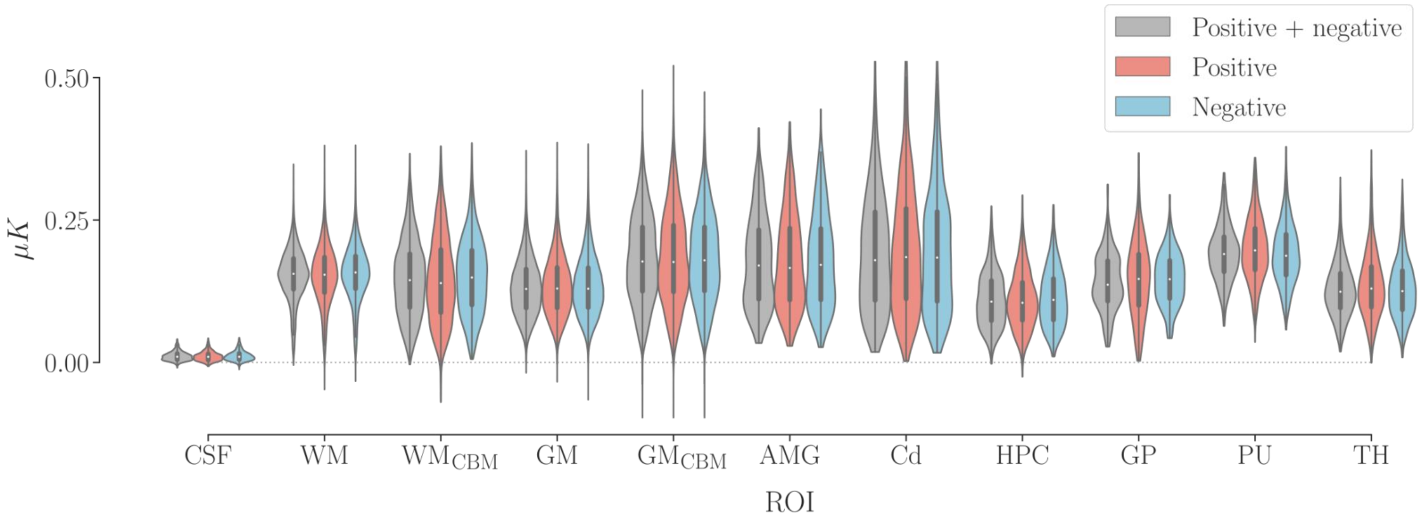


**Fig. S3, Effect of considering only one repetition on each ROI $\mu K$ distribution, without correcting for the cross-terms with imaging gradients.** Distributions of values for each ROI from MNI-space CTI analysis maps, for maps derived from data corresponding to i) two repetitions of the CTI protocol, by alternating the polarity of the diffusion gradients allowing correction for the cross-terms with the imaging gradients ("Positive + negative", grey), ii) one repetition, with "positive" polarity of the diffusion gradients ("Positive", red), iii) one repetition, with "negative" polarity of the diffusion gradients ("Negative", blue). Abbreviations: CSF: Cerebrospinal fluid in lateral ventricles; WM: White Matter; $WM_{CBM}$: Cerebellar White Matter; GM: Grey Matter; $GM_{CBM}$: Cerebellar Grey Matter; AMG: Amygdala; Cd: Caudate; HPC: Hippocampus; GP: Globus Pallidus; PU: Putamen; TH: Thalamus.
